# BioSecBench-Function: A Verifiable Benchmark for Reasoning about Biological Function from Experimental Data

**DOI:** 10.64898/2026.09.03.749010

**Authors:** Dianzhuo Wang, Qian Xu, Arjun Banerjee, Rishi Jain, Aakarsh Vermani, Jonathan Felix, Gage Moreno, Faisal AlZaben, Nicholas Keul, Evan Seeyave, Harmon Bhasin

## Abstract

Inferring biological function from experimental data is central to understanding emerging pathogens and developing effective countermeasures, yet interpreting these data remains slow and expert-intensive. AI agents could help accelerate this process by reasoning across sequence, structural, and biophysical evidence. We present BioSecBench-Function, a verifiable benchmark for recovering biosecurity-relevant function from real biological data. The benchmark comprises 111 evaluations built from published datasets and graded deterministically against ground truth. We organize evaluations along two dimensions: threat axis (spanning seven biosecurity-relevant question types) and biological question (indicating whether the solution depends primarily on sequence, structure, or biophysical assay data). Across 7,326 runs from twenty-two model-harness configurations, Opus 5 under Claude Code led on endpoint pass rate at 50.3%, and Grok 4.6 under Grok Build led on overall pass rate at 44.1% when refusals counted as failures. Performance varied substantially across both model-harness configurations and task categories. Refusal rates differed sharply by provider, and cost was a poor predictor of accuracy: several configurations exceeded 40% pass rate at low cost. BioSecBench-Function provides a standard for measuring whether agents can be trusted to interpret what a new pathogen or variant does when the next outbreak arrives.

## Introduction

Recovering biological function from experimental data is crucial for understanding pathogen behavior and developing counter-measures [1]. Advances in sequencing and experimental assays produce this data faster and at greater scale [2]. The bottleneck is now interpretation: researchers must integrate evidence across modalities and determine which measurements are relevant, reliable, and sufficient to support a functional conclusion.

This matters most during an outbreak, when interpreting a variant’s function determines which variants to prioritize. During the SARS-CoV-2 pandemic, high-throughput mutational scanning of the viral receptor-binding domain mapped how mutations affected folding and ACE2 binding [3, 4]. Researchers then built models on these measurements to predict which variants would be more infectious [5, 6, 7] and which antibody cocktails were most vulnerable to escape [8]. This work was slow and expert-intensive; doing it faster would shorten the path from a new variant to a response.

Agents have shown increasingly strong performance on biological reasoning and computational tasks [9], raising the possibility that they could accelerate functional interpretation. Yet no benchmark measures this directly. Existing execution-based

## Benchmark construction

Each evaluation is built around a published study or dataset spanning modalities such as deep mutational scanning[3], Tite-Seq[17], SPR[18], X-ray crystallography[19], or free-energy calculations[20]. The agent is asked to recover a specific functional quantity from the data provided. The 111 evaluations were constructed by experimental and computational biologists and reviewed before inclusion; see Methods for details of the deterministic grading pipeline.

### Evaluation inventory

We annotate each evaluation along two dimensions (Figure 1).

**Figure 1:**
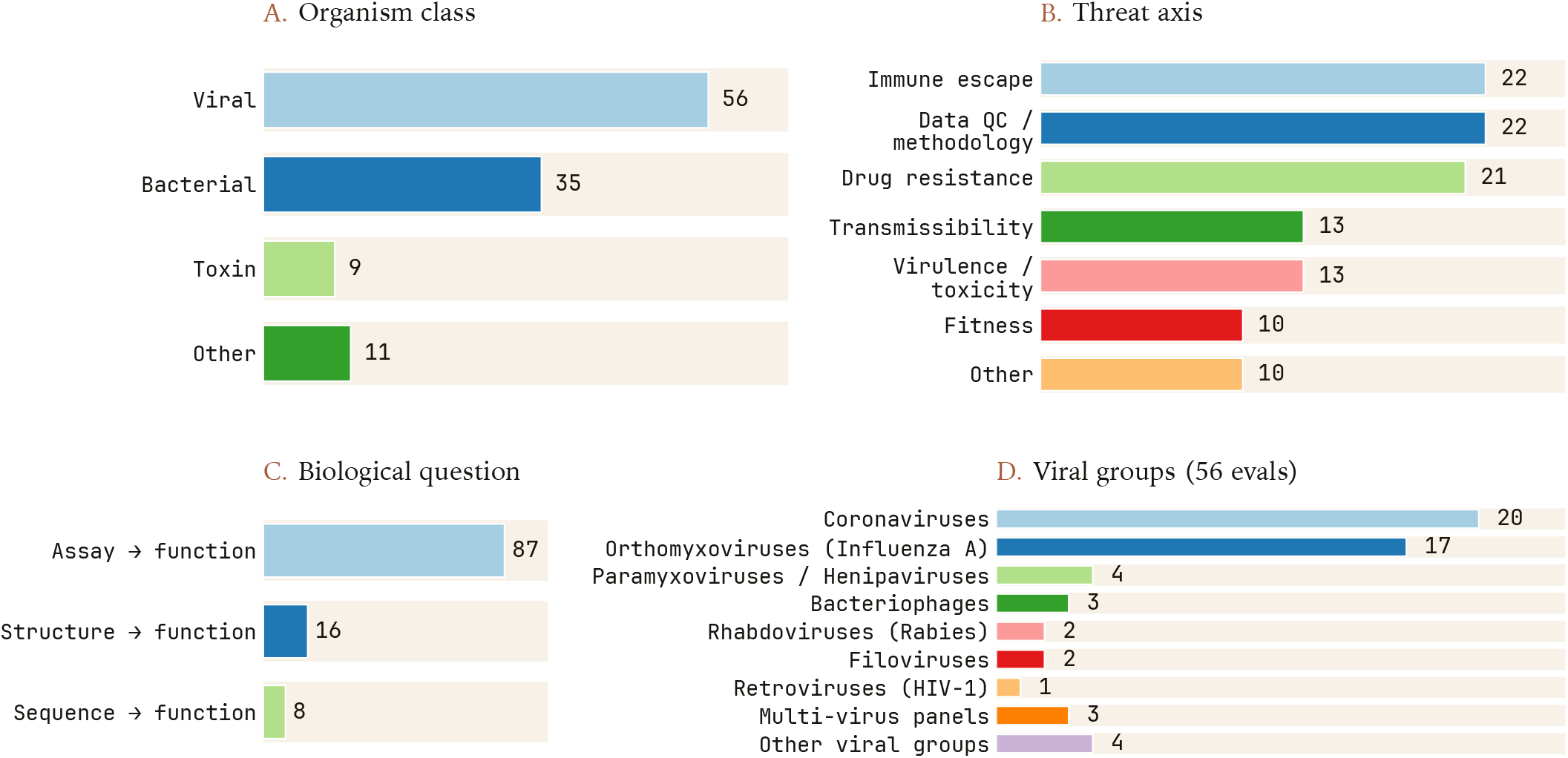
Composition of the 111 BioSecBench-Function evaluations. (A) Organism class. (B) Threat axis. (C) Biological question. (D) Viral groups for the 56 viral evaluations.

The first is the **threat axis**, the biosecurity-relevant function being measured: *transmissibility, immune escape, virulence/toxicity*, benchmarks test whether agents resolve software engineering or computational-biology tasks [10, 11, 9, 12, 13], while a separate line of work evaluates static machine learning predictors on curated fitness-landscape datasets [14, 15, 16]. Neither gives an agent biological data from viruses, bacteria, or toxins and asks it to reason its way, without a predetermined pipeline, to the correct functional conclusion.

BioSecBench-Function is a verifiable benchmark that measures whether AI agents can recover biosecurity-relevant biological function from experimental data, including assay readouts, structures, and protein sequences. Each of its 111 evaluations is built from published data and graded deterministically against ground truth derived from those same data.

Current agents remain far from reliable on this task. Across twenty-two model-harness configurations, the strongest configuration achieved an endpoint pass rate of about 50%, while the average passed little more than a third. Performance also varied by task category: pass rates differed by more than 2× across threat axes and by a comparable margin across biological-question modalities. When refusals were scored as failures, the threat-axis gap narrowed. We release BioSecBench-Function as a standard for measuring whether agents can be trusted to characterize emerging threats when the next outbreak arrives.

*drug resistance, fitness, data QC/methodology*, or *other*.

The second is the **biological question**, whether function is inferred from sequence, structure, or biophysical assay data: *sequence→function, structure→function*, or *biophysical assay→function*.

The benchmark’s coverage reflects what published data are available. Of the 56 viral evaluations, coronaviruses (20, almost all SARS-CoV-2) and influenza A (17) account for 37. The remaining 19 are spread across paramyxoviruses/henipaviruses, bacteriophages, filoviruses, rhabdoviruses (rabies), retroviruses (HIV-1), multi-virus panels, and one-off entries folded into “Other viral groups” (Figure 1D). We use “viral group” to represent these heterogeneous taxonomic and biological groupings. By biological question, 87 of 111 evaluations use biophysical assay data, 16 require structural interpretation, and 8 require sequence inference (Figure 1C).

## Results

### No configuration exceeds *∼* 50% mean endpoint pass rate

Across twenty-two model-harness configurations, endpoint pass rates ranged from 25.4% (GPT-5.6 Luna under PI) to 50.3% (Opus 5 under Claude Code) and averaged 37.6% (Figure 2B). We define endpoint pass rate as the fraction of gradable attempts that are correct, excluding refusals from the denominator. Each configuration pairs one model with one inference harness, the agent scaffold (Claude Code, PI, Mini-SWE-Agent, OpenAI Codex, or Grok Build) that gives the model its tools and prompt structure. The ranking tracks provider closely: the top three configurations are all Anthropic and the bottom six all OpenAI, even though each cluster spans multiple models and harnesses.

**Figure 2:**
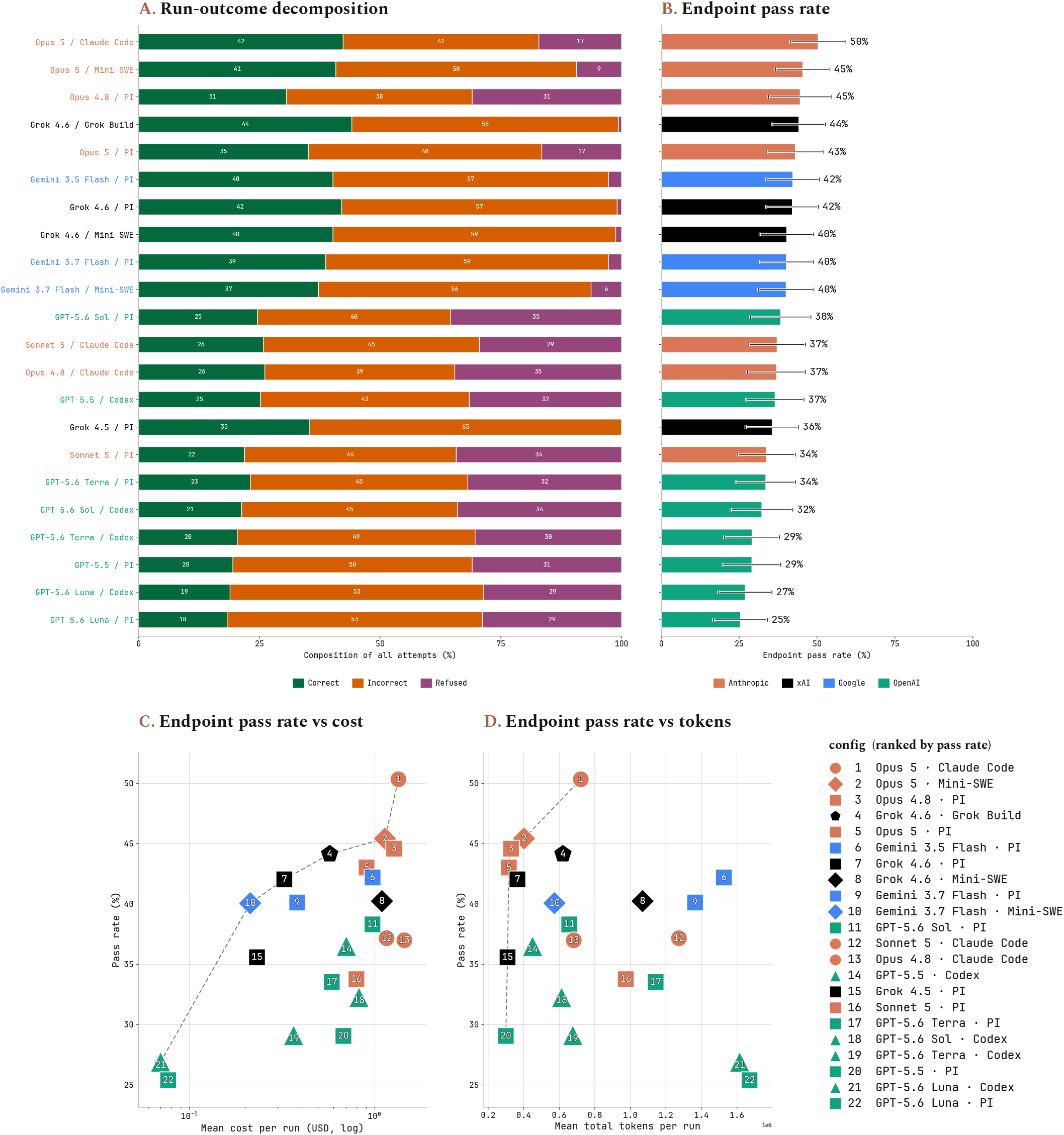
Topline BioSecBench-Function performance, cost, and tokens across twenty-two model × harness configurations. (A) Run-outcome decomposition: every attempt is correct, incorrect (a wrong answer, a timeout, or an ungradable answer), or refused, as a share of all attempts. Endpoint pass rate per configuration, computed as the mean per-evaluation fraction of gradable attempts that were correct, with refusals excluded; error bars are 95% *t* intervals over evaluations. Endpoint pass rate versus (C) mean cost per run in USD (log scale) and (D) mean total tokens per run, both counted under standard input, output, and cached-token conditions. There is one marker per model × harness configuration, colored by provider and shaped by harness, numbered by pass-rate rank (key at right). Resource values are averaged over non-refused, completed runs only. The dashed line is the Pareto frontier: configurations not dominated by another configuration with both a lower resource cost and a higher pass rate.

Endpoint pass rate, rather than an overall pass rate that scores refusals as failures, is our primary metric for two reasons. First, capability can only be demonstrated on questions a model attempts; scoring refusals as failures would penalize a model’s safeguards rather than its capability. Second, not every refusal is a capability failure: several tasks come from dual-use work, where refusing is sometimes correct. Appendix B reports pass rates under the stricter, refusal-penalized convention as a robustness check. Under that convention, Opus 5 under Claude Code drops to second (42.3%) while Grok 4.6 under Grok Build rises to first (44.1%).

Refusal rates varied sharply by provider (Figure 2A): OpenAI configurations refused 31.4% of attempts and Anthropic 24.6%, against 3.9% for Google and 0.68% for xAI. Appendix A analyzes refusal types and their problem domains.

### Higher endpoint pass rates cost more but do not require more tokens

Higher endpoint pass rates broadly track higher cost, but with diminishing returns: several of the strongest configurations cost far less than the top scorer (Figure 2C). Opus 5 under Claude Code, the strongest configuration at 50.3%, is also among the most expensive at $1.34 per run. The pass rate drops off immediately down the frontier: Opus 5 under Mini-SWE-Agent scores 4.9 percentage points lower despite costing only $0.21 less per run. Grok 4.6 under xAI’s native Grok Build harness scores 44.1% at $0.57 per run, roughly half the cost of the three configurations ranked above it. The frontier then continues through Grok 4.6 under the PI harness (42.0% at $0.33 per run) and Gemini 3.7 Flash under Mini-SWE-Agent (40.1% at $0.21 per run).

The cheapest configurations overall, GPT-5.6 Luna under PI and Codex ($0.07–0.08 per run), are also among the weakest (25– 27%). Optimizing for cost alone therefore risks a steep drop in endpoint pass rate.

Token count tells a different story: the highest pass rates do not require the most tokens. Opus 5 and Opus 4.8 under PI reach 43.0% and 44.6% at only *∼*320K tokens per run (Figure 2D), putting both in the top quartile of endpoint pass rate while sitting in the bottom quartile of token use. Even Opus 5 under Claude Code, the strongest configuration, uses fewer tokens than eight of the other twenty-one configurations tested. Several of the heaviest token users, meanwhile, cluster well below the frontier. Cost and tokens are not interchangeable either: GPT-5.6 Luna uses the most tokens of any configuration yet is also the cheapest, since token pricing differs by provider and model.

### Performance varies by threat axis, organism, and data modality

Endpoint pass rate varied more than 2× across the named threat axes (Figure 3A): transmissibility was the easiest at 64.9%, followed by fitness (46.2%) and drug resistance (42.8%), with virulence/toxicity (27.9%) and immune escape (30.7%) the hardest. Data QC/methodology, a non-threat category kept separate from the five threat axes, scored 37.4%. This gap narrows under the refusal-penalized overall pass rate (Appendix B): transmissibility falls to 38.1% and virulence/toxicity, still the hardest named axis, rises to 24.8%, roughly a 1.5× spread rather than 2×.

**Figure 3:**
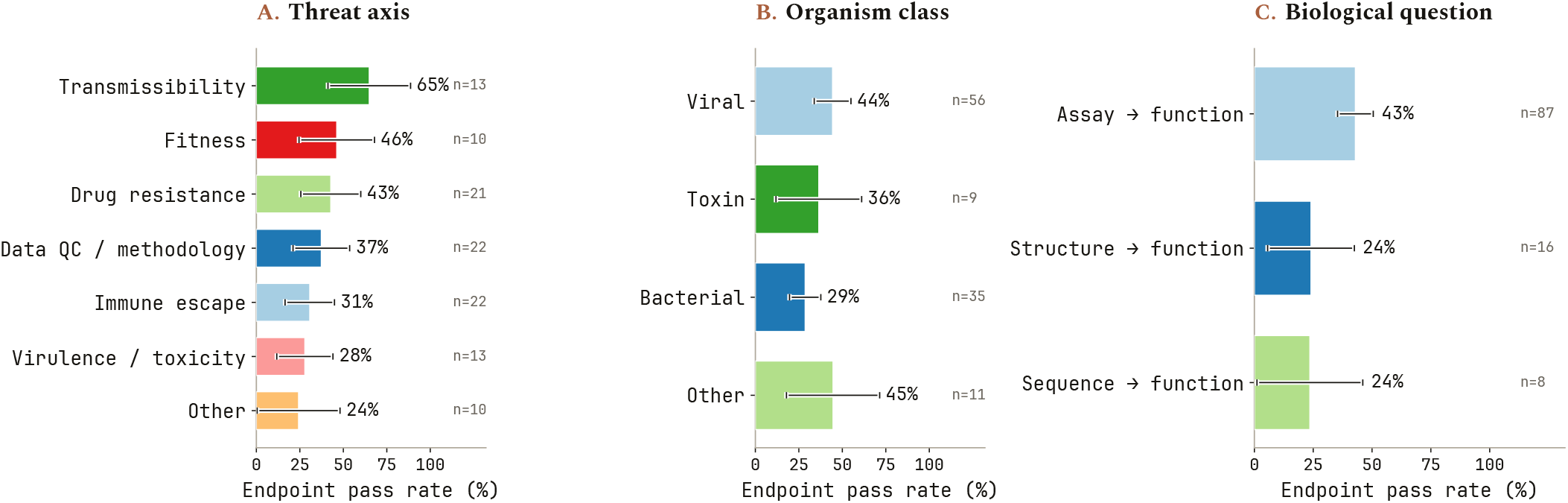
BioSecBench-Function performance broken down by evaluation attributes. Endpoint pass rate by (A) threat axis, (B) organism class, and biological question, pooled across all twenty-two configurations. Each evaluation was reduced to one pass-rate estimate; n is the number of evaluations, and cells with fewer than 5 evaluations are omitted. Error bars are 95% *t* intervals over evaluations.

The axis ordering held across every configuration (Figure 4): transmissibility ranked among the two easiest for all 22. Fitness narrowly took the single easiest spot in only two, GPT-5.5 under PI and GPT-5.6 Luna under Codex. Virulence/toxicity ranked among the two hardest for 15 of the 22.

**Figure 4:**
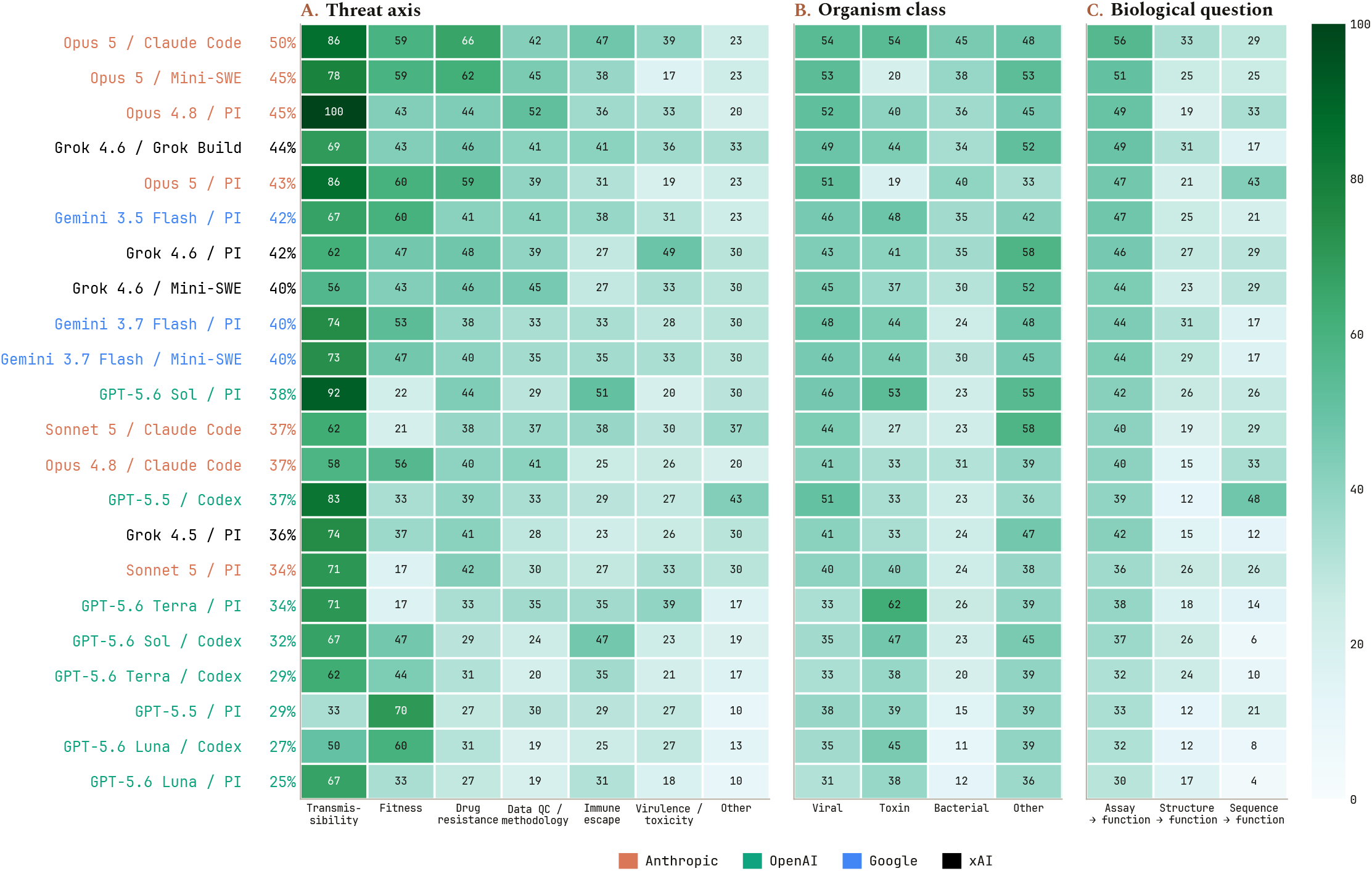
Endpoint pass rate by configuration and threat axis, organism class, and biological question. Each cell is the eval-level mean endpoint pass rate for one model-harness configuration on one category; darker is higher. Rows are configurations ordered by endpoint pass rate (shown beside each label; strongest at top) and colored by provider. Columns are (A) threat axes, (B) organism class, and (C) biological question, ordered by endpoint pass rate (easiest at left).

### Failure patterns reflect incorrect scientific judgment

To better understand where agents went wrong, we conducted trajectory analysis and found instances where agents lacked scientific judgment. We illustrate two types of error below: first, agents forced conclusions from inconclusive data; second, they skipped well-established QC steps when interpreting assays.

The first pattern appears in a task asking whether the SARS-CoV-2 JN.1 variant escapes an antibody candidate strongly enough to reject it. The agent receives conflicting evidence: three monovalent measurements show weak binding, a multivalent avidity assay shows strong binding. Because the two formats probe different interaction regimes, the data cannot distinguish true escape from an assay-format artifact, so the correct call is to defer until a confirmatory experiment is run, which ultimately approves the candidate. Nearly every agent rejected it instead: Opus 5 scored 0/8, Opus 4.8 scored 0/6, and every Gemini and Grok configuration scored zero. Even recognizing the gap did not change the call: Opus 5 often named the missing experiment, then rejected the candidate. Only GPT-5.6 Luna consistently deferred (5/6).

The same error appears in a different form when agents invent a rule unsupported by the data. In this evaluation, each protein domain’s folding stability is measured twice, once with Organism class and biological question show the same wide spread as the threat axes. Among the named organism classes (Figure 3B), viral evaluations scored highest (44.4%), followed by toxin (36.4%), with bacterial the lowest of the well-populated classes (28.6%). By biological question (Figure 3C), agents recovered function far more reliably from biophysical assay data (42.9%) than from structure (24.1%) or sequence (23.6%) alone. trypsin and once with chymotrypsin. Because both proteases are equally valid readouts, averaging them gives the right answer. Instead of averaging, a failing Opus 4.8 run took the lower value for each domain, justifying it with a rule it invented: that one protease was “the limiting protease in the assay.”

The second pattern, skipping a necessary quality check, appears in a carbapenemase mutation scan. Carbapenemase is the enzyme bacteria use to break down a last-resort class of antibiotics. Each resistance measurement comes with an error bar, and those bars vary more than a hundredfold: values near complete resistance or complete sensitivity are far noisier than those in the middle. Getting the call right means judging each measurement against its own error bar. Agents that weighed each measurement against its own error bar scored well (Opus 5 9/9, GPT-5.6 Sol 6/6), while those that applied one fixed cut-off mistook noise for resistance and scored 0/6 (GPT-5.5, GPT-5.6 Luna).

The same failure to check data quality appears in a binding free-energy simulation between a bacterial toxin and its cognate immunity protein. The simulation needs an initial equilibration period before its output is reliable. Agents that discarded the unequilibrated opening recovered the correct binding free energy (GPT-5.6 Sol and GPT-5.6 Terra 6/6, Opus 5 9/9); those that averaged over the whole trajectory overestimated it and failed (Opus 4.8 and GPT-5.6 Luna 2/6).

## Discussion

BioSecBench-Function measures whether AI agents can recover biosecurity-relevant function from experimental data. Across every configuration we tested, they cannot do so reliably: only Opus 5 under Claude Code cleared a 50% endpoint pass rate (50.3%), and it falls to 42.3% once refusals are scored as failures (Appendix B). This mirrors our finding in BioSecBench-Surveillance [21] despite the two benchmarks testing distinct capabilities, and the case studies suggest these failures stem in part from a lack of scientific judgment.

Performance also varied widely across threat axes, and part of that gap is data modality. Every one of the twenty-two configurations scored higher on transmissibility than on virulence/toxicity or immune escape, regardless of model, harness, or provider. Transmissibility is built almost entirely from biophysical assays (12 of 13 evaluations), the modality agents handle best (42.9%), whereas virulence/toxicity carries the highest share of structure-based tasks of any named axis (31%), the modality agents handle worst (24.1%).

One confounder, refusals, complicates any capability claim drawn from these numbers, because a model cannot demon-strate competence on a task it never attempts and the set of attempted tasks varies widely across configurations. The same model can swing on this alone: Opus 4.8 scores 100% on transmissibility under PI but 58% under Claude Code, because the higher-scoring harness refuses tasks the other attempts and fails (Appendix B). This is enough to change the headline: under the refusal-penalized overall pass rate, Grok 4.6 under Grok Build overtakes Opus 5 under Claude Code, 44.1% versus 42.3%, because Opus 5 refuses 17% of its attempts against Grok Build’s 0.6%. Because the direction and magnitude vary by configuration, refusals confound capability rather than bias it in a correctable way, and these benchmarks should be read alongside dedicated refusal benchmarks.

Several limitations bound how far these results generalize. Evaluations are unevenly distributed across threat axes and biological-question categories, and each category label comes from mapping a task onto a fixed taxonomy. Subgroup sizes are small once results are split by axis or organism class, so per-category pass rates carry high variance and should be read with caution. The benchmark also includes only tasks with a single deterministically gradable answer, leaving open-ended functional judgments outside its scope. Tasks were curated from failure modes observed in agents, so a pass rate reflects performance on a deliberately hard subset of an axis, and cross-axis comparisons should be read the same way. Finally, our trajectory analysis is partial: it rests on manual inspection of selected runs, and a systematic pass could separate agents that never consider the right interpretation from those that consider and reject it, cluster recurring failure rationales, and quantify how the harness shapes exploration and commitment.

Detection is only the beginning of biosecurity response; what ultimately matters is how newly observed variants, toxins, and pathogens are interpreted to inform countermeasures. BioSecBench-Function measures that step directly and shows that current agents remain unreliable at it even when the relevant data are available and the answer is objectively gradable. As biological data generation accelerates, strengthening the scientific judgment these tasks demand will be critical to making agents useful in biosecurity.

## Methods

### Benchmark composition and data

Each evaluation is a single definition file (task prompt, grader, metadata tags, and pointers to input data), plus a structured notes field documenting what the evaluation tests and why, for example what property of the underlying data makes an easy-looking shortcut wrong. The 111 evaluations were constructed by a team of eight domain experts from published datasets spanning viral, bacterial, toxin, and a small number of other organism systems, and each is documented in a benchmark card recording organism class, pathogen status, source paper, assay modality, and grading rationale.

### Task format, deterministic grading, and refusal classification

Every evaluation is graded by a deterministic grader that allows for many forms and combinations of outputs. For numeric outputs, fields are graded against either a relative or absolute tolerance or an explicit range. For set outputs, grading is done via label overlap against a Jaccard threshold. Categorical or multifield answers are assessed by per-key dictionary match, predicate, or multiple choice. Composite evaluations combine several graders by requiring that every sub-grader passes. Tolerances are set per evaluation from the underlying biology and how the ground truth was derived. A run passes only when every required check passes; missing fields, invalid JSON, and off-schema or unparseable answers count as failures.

API refusals were classified by matching trajectory outputs to known model- and harness-specific refusal signatures, while model-initiated refusals were classified by running an LLM judge over the trajectories. This allowed us to detect both system-level and model-level blocks.

### Agent runs and execution

Each model-harness pair was run three times per evaluation, in an isolated sandbox with no internet access, so an agent cannot look up the source paper, the organism, or the answer and must work from the files staged in the evaluation itself.

A configuration is a model paired with an inference harness, the scaffold (Claude Code, PI, Mini-SWE-Agent, OpenAI Codex, or Grok Build) that gives the model its tools and prompt structure; as in BioSecBench-Surveillance, we treat model and harness as independent axes because the same model can behave very differently across them, in both capability and refusal; Gemini 3.7 Flash and several Anthropic models are each tested under two harnesses here for exactly this reason. Every run’s complete raw trajectory (the conversation, tool calls, and execution outputs) is recorded. We evaluated twenty-two deployed configurations: Opus 5 under Claude Code, Mini-SWE-Agent, and PI; Opus 4.8 and Sonnet 5 under Claude Code and PI; Gemini 3.5 Flash under PI; Gemini 3.7 Flash under PI and Mini-SWE-Agent; GPT-5.5, 5.6-Luna, 5.6-Sol, and 5.6-Terra under PI and OpenAI Codex; and Grok 4.5 under PI and Grok 4.6 under PI, Mini-SWE-Agent, and Grok Build.

All models are run at their maximum reasoning-effort level. This yielded 111 evaluations × twenty-two configurations × 3 trials = 7,326 runs. Per-run cost (USD) and token usage (input, cached-input, and output) were also recorded; Figures 2C,D report these averaged over non-refused, completed runs only, since a refused or errored run does not reflect the resource cost of a full analysis attempt.

### Outcome classification and aggregation

Each run is assigned one of three outcomes, correct, incorrect, or refused, that sum to the run count for every evaluation and configuration. The endpoint pass rate is correct runs over the sum of correct and incorrect runs (excluding refusals); timeouts and errored runs count as incorrect. Refusals are classified into API-level blocks or model-reasoning-level blocks. Some models and harnesses enable tool-call blocks as well; for BioSecBench-Function, a mid-trajectory block that did not end the run is recorded but not counted as a refusal, since the run continued to a gradable answer regardless. Uncertainty intervals are 95% Student-*t* intervals computed across per-evaluation pass rates, with the evaluation as the sampling unit.

## Data Availability

Consistent with accepted biosecurity research practice, the full 111-evaluation set is held under restricted access. A public subset of evaluations and benchmark construction code is available at https://github.com/latchbio/biosecbench-function.

## Appendix A: Refusal Analysis

The provider gap in refusal rate (Figure 2A) masks a sharper difference in how refusals arise. Every Anthropic refusal is an API-level block, with no model-initiated declines at all. The same holds nearly true for OpenAI and Google models, where essentially all refusals (99.4% and 97.4%, respectively) are API-level rejections rather than model-initiated declines. In contrast, xAI’s Grok configurations show almost no refusals (9 of 1,332 runs), and all nine are model-initiated; we observed no API-level blocks for xAI. Thus, for three of the four providers, refusals are dominated by provider-side filtering, whereas for xAI they arise almost exclusively from the model itself (Figure 5).

**Figure 5:**
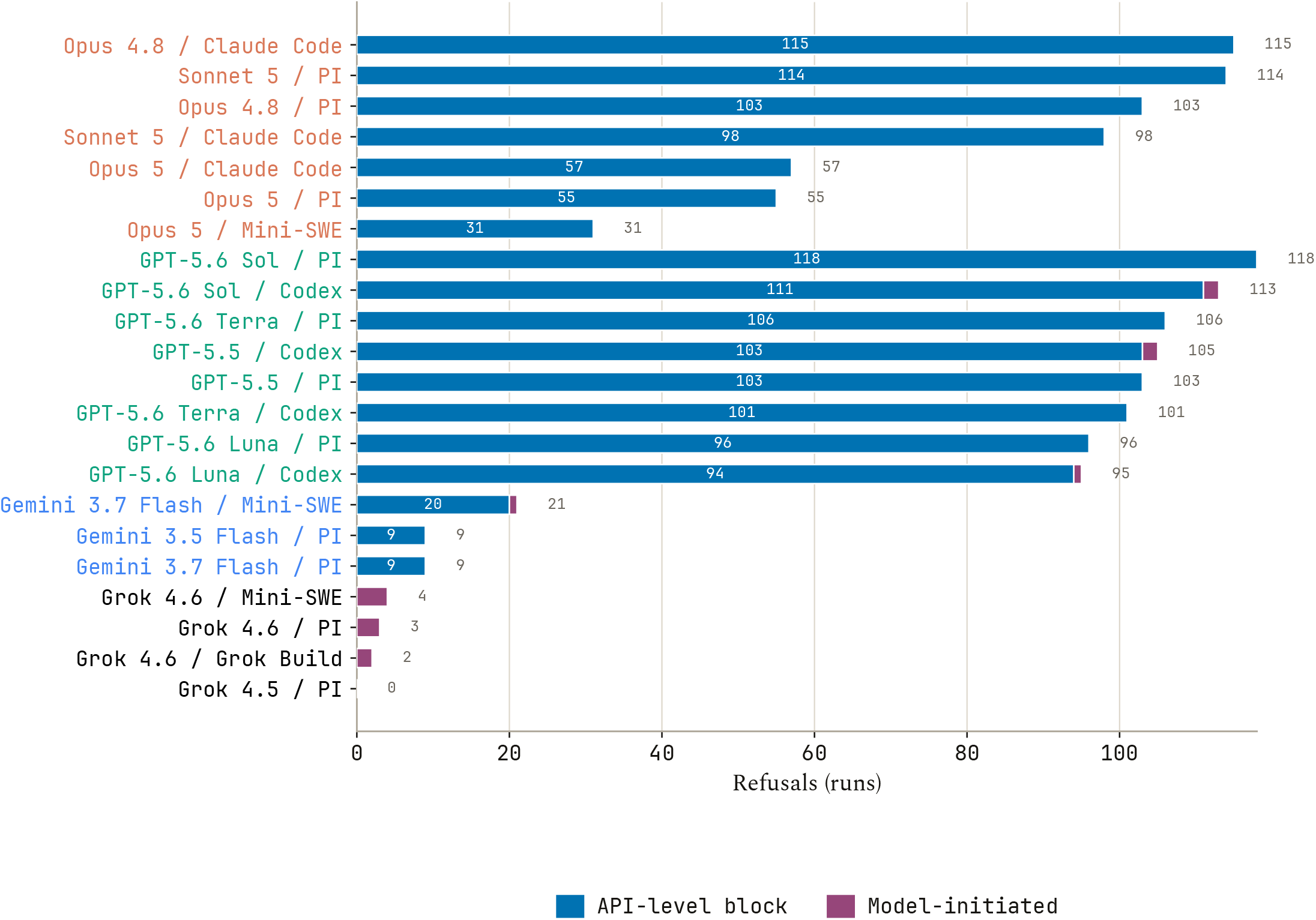
Refusal types across model-harness combinations. Each horizontal bar is one model-harness configuration. The total length of a bar is the number of runs that ended in a refusal, with the count indicated to the right of the bar. Bars are decomposed into API-level blocks (the harness or provider infrastructure rejects the request or response) and model-initiated refusals (the model itself declines the task in its answer). Ordering and coloring are based on providers; ordering inside provider-groups is by the number of refusals.

One caveat for refusals is that there are, very rarely, additional tool-level refusals that fire when the API deems the inputs or outputs of tool calls to be potentially hazardous. For some providers, such as xAI, tool-level refusals are non-terminal, and the model can still continue with reasoning despite the block. In this case, we do not count this as a terminal refusal, since the model can still succeed and pass the task despite this occurring. However, if tool-level refusals do end the trajectory, we do count this as a terminal refusal.

As is common biosecurity practice, the exact breakdown of refusal behavior along threat axis, organism class, or viral group is withheld. A full breakdown is available to qualified partners on request.

## Appendix B: Overall Pass Rates

As mentioned in the Discussion, a natural confounder of measuring capabilities of a model in biosecurity is that the safeguards between configurations provide an uneven distribution of questions attempted. The endpoint pass rate allows for an assessment of capability without punishing the model for strong safeguards, since only answered questions are scored; however, this method falls victim to the unequal distribution concern. As such, overall pass rates, which count refusals as failures, are depicted in Figures 6, 7, and 8.

**Figure 6:**
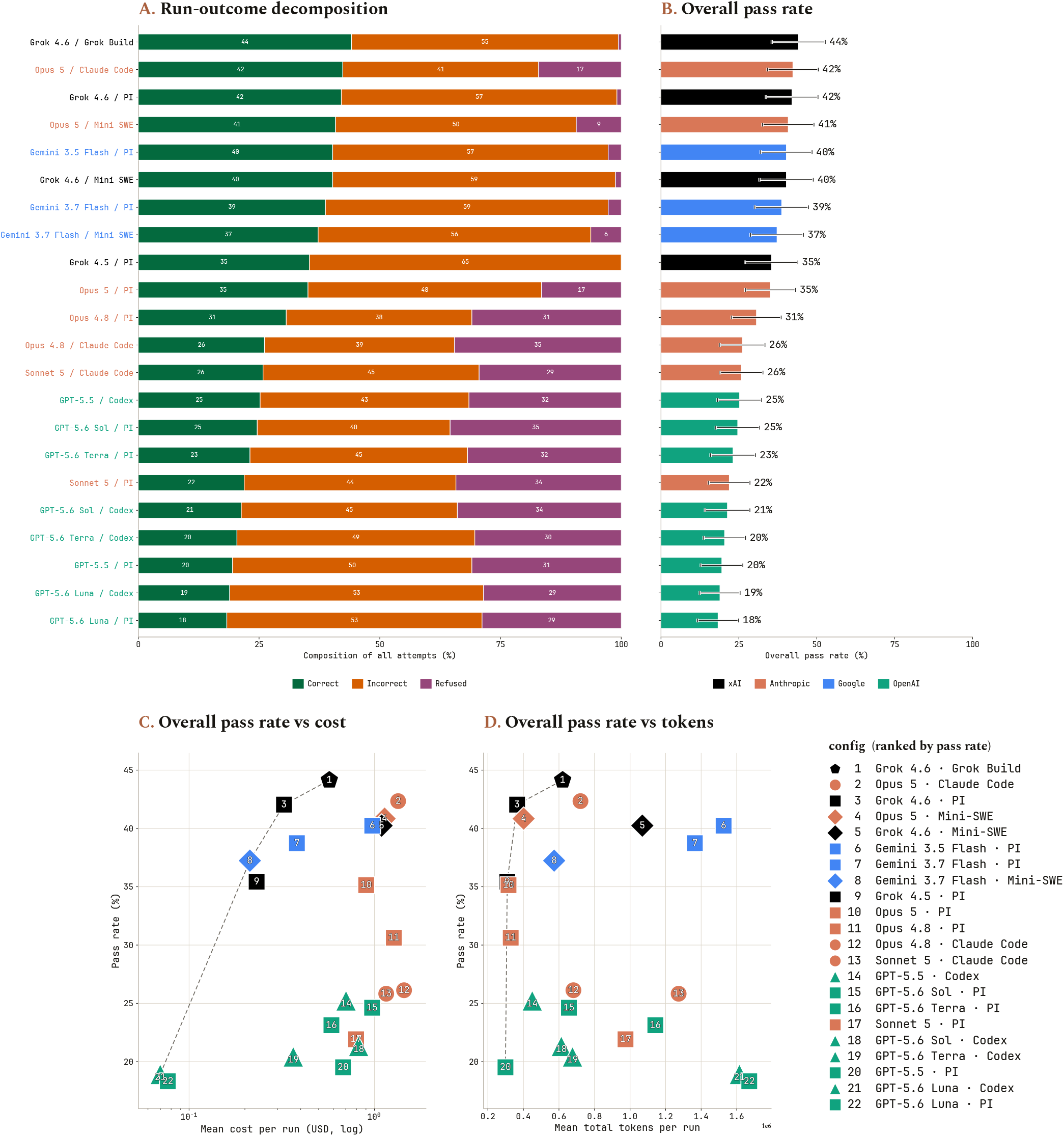
Topline BioSecBench-Function performance under overall pass rate. Companion to Figure 2, with refusals folded into the denominator. (A) Run-outcome decomposition: every attempt is correct, incorrect (a wrong answer, a timeout, or an ungradable answer), or refused, as a share of all attempts. (B) Overall pass rate per configuration, computed as the mean per-evaluation fraction of *all* attempts that were correct (correct / num_runs), so refusals and errored or no-answer runs count as failures; error bars are 95% *t* intervals over evaluations. Overall pass rate versus (C) mean cost per run in USD (log scale) and (D) mean total tokens per run, one marker per model × harness configuration, colored by provider and shaped by harness, numbered by pass-rate rank (key at right). Resource values are averaged over non-refused, completed runs only. The dashed line is the Pareto frontier.

**Figure 7:**
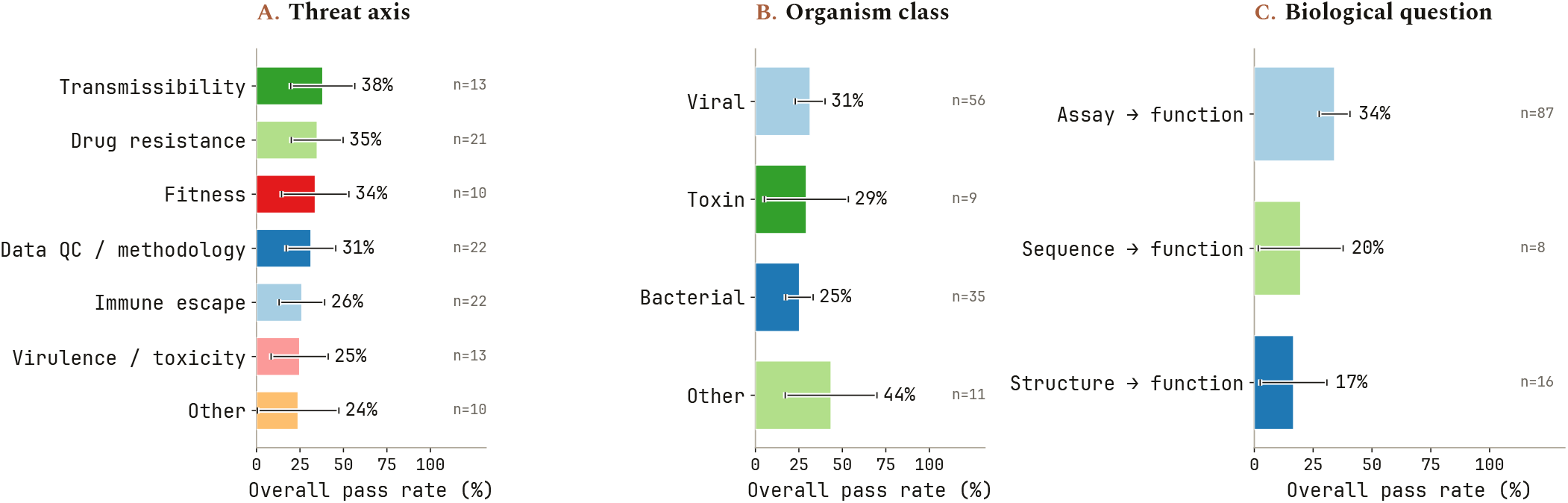
BioSecBench-Function overall pass rate broken down by evaluation attributes. Companion to Figure 3, with refusals folded into the denominator. Overall pass rate (correct / num_runs) by (A) threat axis, (B) organism class, and (C) biological question, pooled across all twenty-two configurations. Each evaluation was reduced to one pass-rate estimate; n is the number of evaluations, and cells with fewer than 5 evaluations are omitted. Error bars are 95% *t* intervals over evaluations.

**Figure 8:**
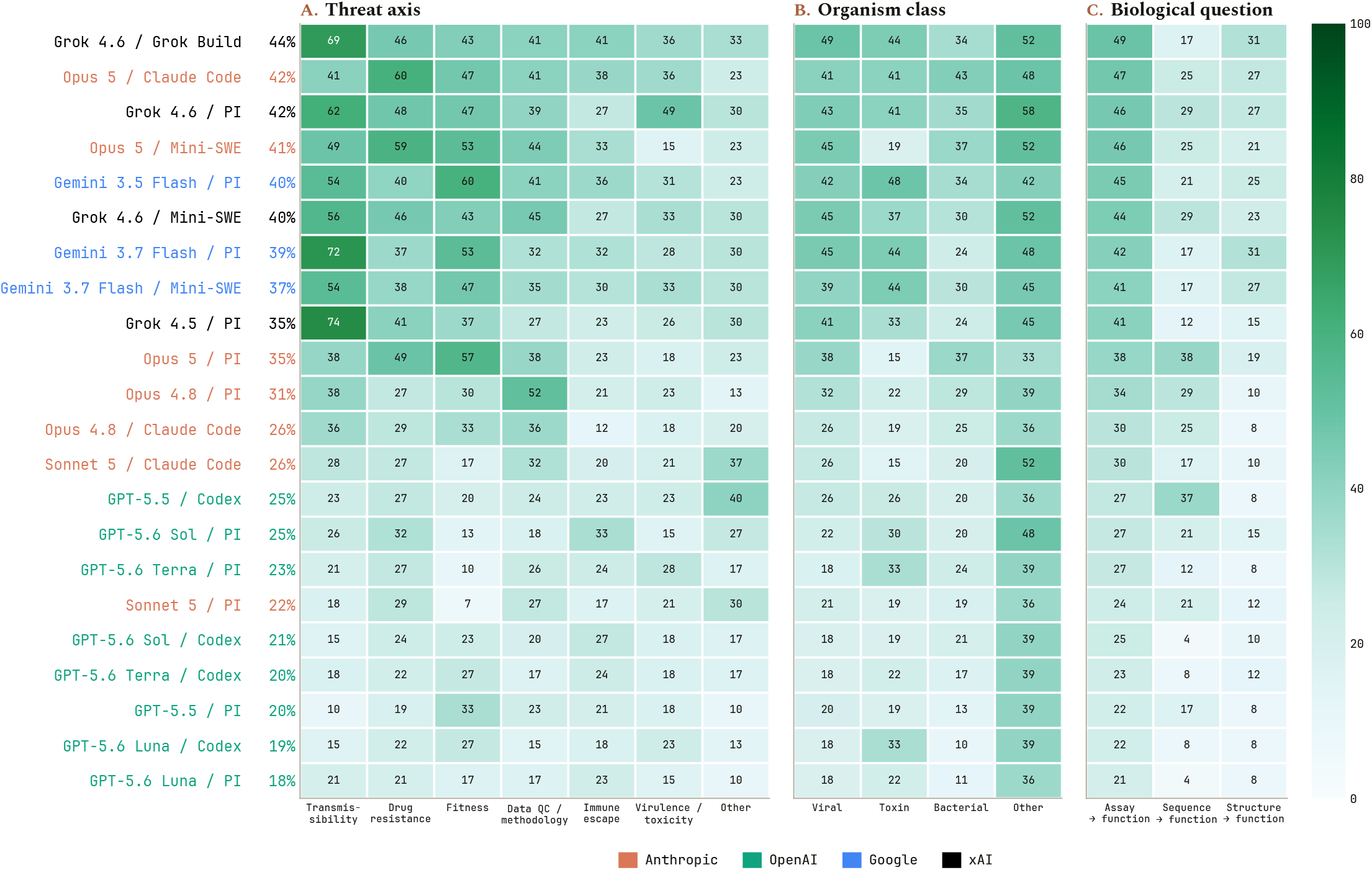
Overall pass rate by configuration and threat axis, organism class, and biological question. Companion to Figure 4, with refusals folded into the denominator. Each cell is the eval-level mean overall pass rate (correct / num_runs) for one model-harness configuration on one category; darker is higher. Rows are configurations ordered by overall pass rate (shown beside each label; strongest at top) and colored by provider. Columns are (A) threat axes, (B) organism class, and (C) biological question, ordered by overall pass rate (easiest at left).

The penalty this convention imposes is not a uniform shift: it averages 7.0 percentage points across configurations but ranges from zero to 14.0, so the ranking itself changes. In Figure 6, Grok 4.6 under Grok Build rises from fourth to first, while Opus 4.8 under PI falls from third to eleventh. Figure 7 shows the corresponding compression of the category gaps, with the spread across named threat axes narrowing from roughly 2× to 1.5×. Figure 8 repeats the per-configuration heatmap under the same convention, where the harness-driven differences noted in the Discussion largely disappear.

Figure 9 puts both conventions on the same axes for every configuration. The dashed line is *y* = *x*, where overall and endpoint pass rate coincide because the configuration never refuses or errors; the vertical drop of a point below that line is exactly the share of its attempts lost to refusals, timeouts, or no-answer runs. Configurations cluster well below the diagonal by similar amounts within a provider, consistent with refusal behavior being driven mostly by provider-level filtering (Appendix A) rather than by each configuration’s own capability.

**Figure 9:**
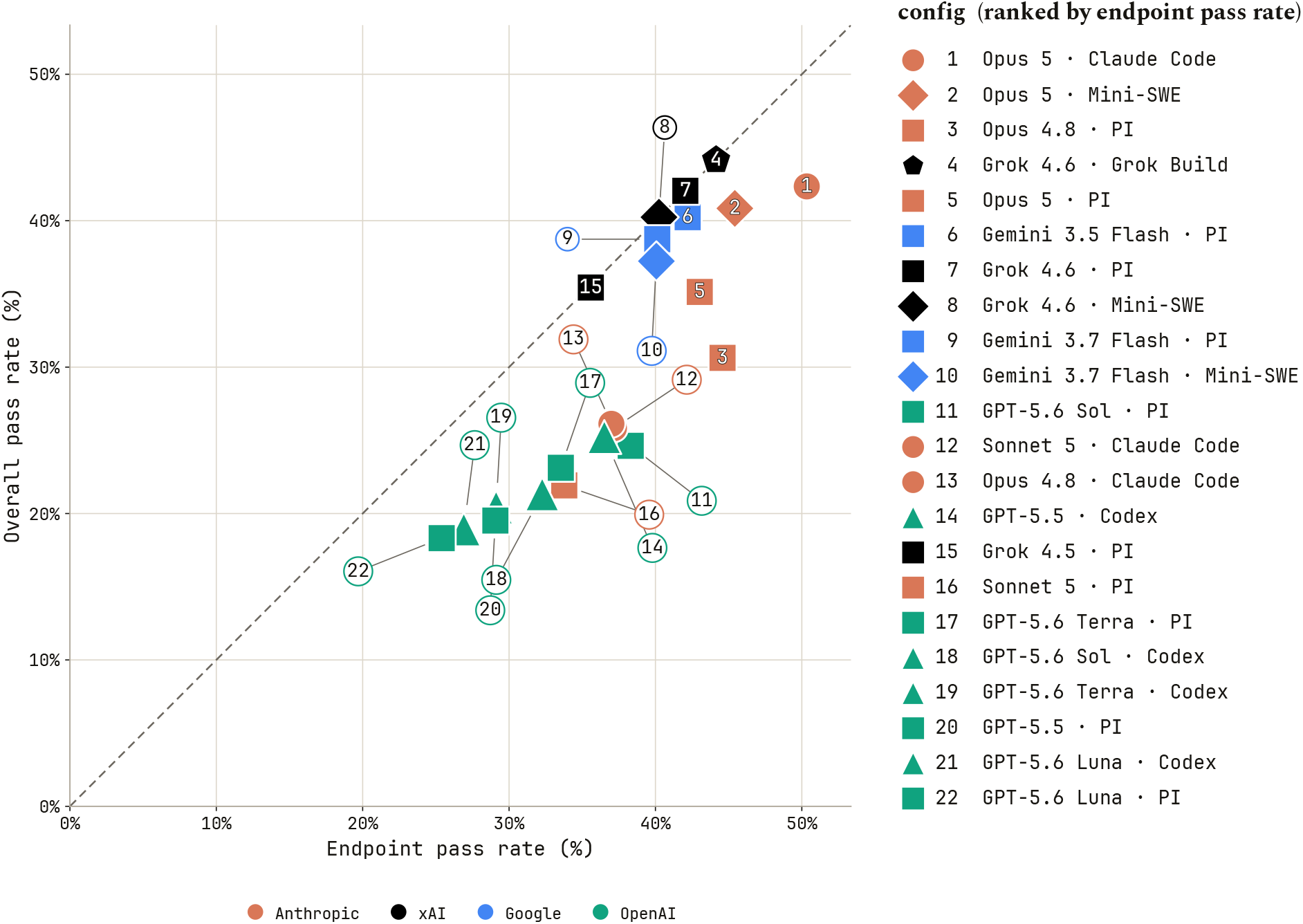
Overall pass rate versus endpoint pass rate, per configuration. Each point is one model × harness configuration at (endpoint pass rate, overall pass rate), colored by provider and shaped by harness, numbered by endpoint pass-rate rank (key at right, shared with Figure 2). The dashed *y* = *x* line marks zero penalty from refusals, timeouts, or no-answer runs; the vertical distance of a point below it is exactly that penalty.

